# in vivo Reprogramming of NG2 Glia Improves Bladder Function After Spinal Cord Injury

**DOI:** 10.1101/2025.07.28.667292

**Authors:** Wenjiao Tai, Junkui Shang, Peiqi Zhao, Wei Li, Tianjin Shen, Xiaoling Zhong, Yuhua Zou, Bo Chen, Chun-Li Zhang

## Abstract

Neurogenic bladder is a debilitating consequence of spinal cord injury (SCI), with few treatment options that restore voluntary voiding. Here, we show that SOX2-mediated in vivo reprogramming of NG2 glia improves bladder function in a clinically relevant mouse model of contusive SCI. NG2 glia reprogramming induces adult neurogenesis, reduces glial scarring, and significantly improves urinary performance, as measured by voiding assays and conscious cystometry. Functional recovery correlates positively with neurogenesis and negatively with glial scarring. These findings demonstrate that SOX2-mediated glial reprogramming promotes autonomic repair and offers a regenerative strategy for neurogenic bladder after SCI.

## INTRODUCTION

Spinal cord injury (SCI) is a devastating neurological condition that leads to a spectrum of motor, sensory, and autonomic impairments ^1,2^. Among these, neurogenic bladder dysfunction is one of the most prevalent and burdensome complications, affecting more than 80% of individuals with SCI ^3-7^. This condition arises from the disruption of descending neural pathways between the brain and spinal cord, resulting in loss of voluntary bladder control, urinary retention, incontinence, and a heightened risk of urinary tract infections. These symptoms not only carry significant medical consequences but also severely diminish quality of life ^7,8^.

Current treatments--such as intermittent catheterization, timed voiding, pelvic-floor exercises, and pharmacologic interventions like anticholinergics or botulinum toxin--are primarily palliative. While they provide partial symptom control, they do not replace lost neurons or restore brain– spinal communication, limiting their long-term efficacy. Thus, there is a pressing need for regenerative approaches that can restore functional neural connectivity and improve autonomic bladder control following SCI.

Over the past decade, substantial progress has been made in identifying therapeutic targets to enhance functional recovery by harnessing spinal neurons following SCI ^9,10^. Nevertheless, SCI still causes progressive neuronal dysfunction and loss ^11,12^, underscoring the critical need to replenish spinal neurons.

Our previous study ^13,14^ demonstrated that in vivo glial reprogramming represents a promising approach to produce new neurons and promote locomotor recovery following cervical dorsal hemisection. Notably, neurons generated from reprogrammed NG2 glia were able to integrate into existing neural circuits, forming synaptic connections with propriospinal neurons as well as neurons in the brainstem and dorsal root ganglia (DRG). Nonetheless, it remains unknown whether a similar SOX2-based strategy can restore bladder function after SCI. In the present study, we investigated SOX2-mediated in vivo reprogramming of NG2 glia in a clinically relevant contusive SCI model and found that this intervention significantly improves bladder function.

## RESULTS

### NG2 glia reprogramming improves bladder function after T13 contusive SCI

NG2 glia exhibit remarkable plasticity and can be efficiently reprogrammed by SOX2 to generate neurons in the adult mouse spinal cord ^13^. To test whether a phospho-mimetic SOX2 mutant--previously shown to enhance reprogramming efficiency in the adult brain ^15^--could achieve similar effects in the spinal cord, we delivered either wild-type or mutant SOX2 via a lentiviral vector driven by the *hNG2* promoter ^13^. To promote neuronal survival and maturation, the neurotrophic factor p75-2 was co-expressed in all conditions ^13^. At 4 weeks post virus-injection (wpv), DCX^+^ immature neurons were robustly detected in both SOX2-expressing groups but not in the p75-2-only control (Fig. S1A-C). However, no substantial difference in neuronal output was observed between wild-type and mutant SOX2. Genetic lineage tracing confirmed that NG2 glia served as the cell of origin for DCX^+^ cells in the mutant SOX2 condition (Fig. S1D, E), and that reprogramming proceeded through an ASCL1^+^ intermediate progenitor stage (Fig. S1F-K). Given the reduced inter-individual variability observed in mutant SOX2-injected mice (Fig. S1B), this construct was used for all subsequent experiments. For simplicity, the mutant SOX2 is hereafter referred to just as SOX2.

To assess the impact of SOX2-mediated NG2 glia reprogramming on bladder dysfunction following SCI, we introduced contusive injuries at the thoracic level 13 (T13) in adult wild-type mice (Fig. 1A). One week post-injury, three viral vectors were injected into the injury penumbra: SOX2/p75-2, p75-2 alone, and GFP alone, with the latter two serving as controls (Fig. 1B). To label newly generated neurons, mice were given BrdU in their drinking water. Bladder function was evaluated using modified spontaneous voiding spot (VSOP) test ^16^ at 11 wpv. Prior to testing, all mice underwent manual bladder expression to ensure complete bladder emptying. Subsequently, each mouse received a subcutaneous injection of saline (50 µL/g body weight) to maintain hydration. Mice were then placed individually in filter paper-lined cages, and voiding patterns were monitored over a 2-hour observation period (Fig. 1C).

**Figure 1.**
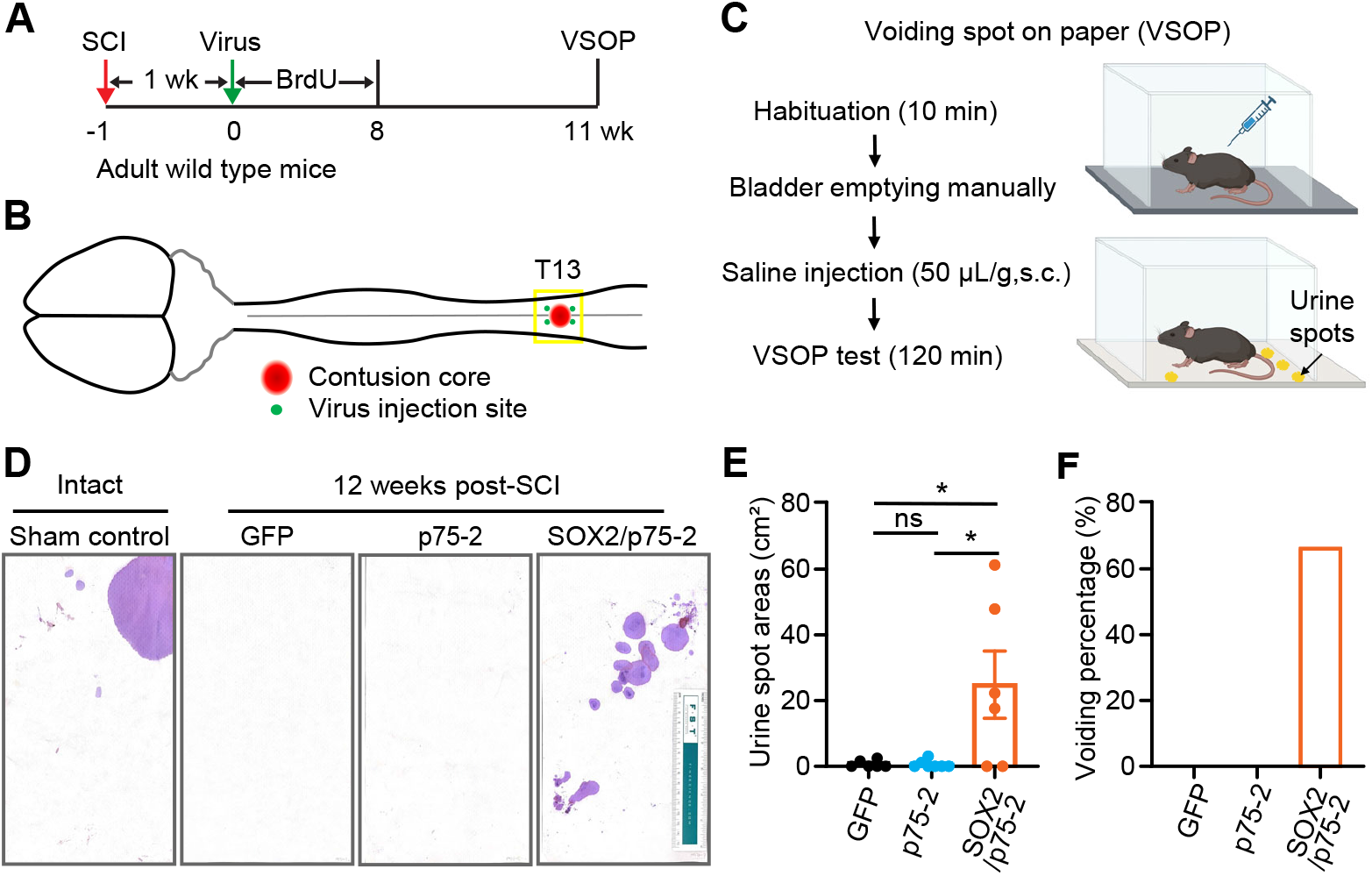
NG2 glia reprogramming improves bladder function after T13 contusive SCI. (A) Experimental scheme. Adult mice received a T13 contusive SCI, followed by viral injection one week later. BrdU was administered in the drinking water to label newly generated neurons. Bladder function was assessed using VSOP tests at 11 weeks post-virus injection (wpv). (B) schematic drawing of T13 contusive SCI and the locations of virus injections. (C) Experimental procedure for VSOP tests. (D) Urine spots for the indicated experimental groups at 11 wpv. (E) Quantification of urine spot areas (mean ± SEM, n = 6-7 mice per group; Ordinary one-way ANOVA: F_(2, 16)_ = 6.077 and *p = 0.0109 for treatment effects; Tukey’s multiple comparisons test: *p = 0.0230 for SOX2/p75-2 vs. GFP, *p = 0.0179 for SOX2/p75-2 vs. p75-2; ns, not significant). (F) Quantification of voiding percentage (mean ± SEM, n = 6-7 mice per group).

VSOP analysis revealed marked differences in voiding patterns across experimental groups (Fig. 1D). Sham-operated mice exhibited normal voiding behavior, with urine spots predominantly localized to the cage corners and minimal central spotting. In contrast, mice treated with either GFP or p75-2 alone showed no visible urine spots. Notably, mice receiving SOX2/p75-2 displayed scattered urine spots on the filter paper, although these were randomly distributed (Fig. 1D). Quantification of total urine spot area revealed an average of approximately 20 cm^2^ in the SOX2/p75-2 group, compared to negligible spotting in the GFP and p75-2 groups (Fig. 1E; F_(2, 16)_ = 6.077 and *p = 0.0109 for treatment effects; *p = 0.0230 for SOX2/p75-2 vs. GFP, *p = 0.0179 for SOX2/p75-2 vs. p75-2). Among the six SOX2/p75-2-treated mice, 67% of them exhibited urine spots, whereas none of the GFP- or p75-2–only mice did (Fig. 1F). Together, these findings suggest that severe T13 contusion SCI abolishes spontaneous voiding within the 2-hour testing window, and that SOX2/p75-2 treatment partially restores spontaneous urinary turnover in injured mice.

### NG2 glia reprogramming reduces neurogenic bladder retention after T13 contusive SCI

The above VSOP analysis revealed that SCI mice treated with SOX2/p75-2 exhibited significantly shorter urination turnover times compared to controls. This improvement may result from two possible mechanisms: (1) SOX2/p75-2 treatment may help restore descending pathways between the brain and sacral voiding center, thereby relieving SCI-induced urinary retention ^17^; and (2) SOX2/p75-2 treatment may enhance local activity within the sacral voiding circuitry, potentially leading to increased bladder contractions and urinary incontinence ^7^.

To explore these mechanisms, we performed cystometry in unrestrained, conscious mice as a terminal experiment at 12 or more wpv (Fig. 2A). In intact mice, continuous saline infusion into the bladder at a constant rate (10 µL/min) gradually fills the bladder, triggering micturition contractions. These contractions appear as spikes in bladder pressure followed by a return to baseline, reflecting bladder emptying. In naive animals, bladder voiding closely correlated with these contractions, typically showing four to five voiding contractions over a 20-minute recording period (Fig. 2B). In contrast, contused mice treated with either p75-2 or GFP displayed frequent, high-frequency, low-amplitude, non-voiding contractions--hallmarks of an overactive bladder (Fig. 2C). While non-voiding contractions persisted in contused mice treated with SOX2/p75-2, this group exhibited improved efficiency of voiding-associated contractions (Fig. 2D). Although the total number of bladder contractions was comparable across all groups (Fig. 2E; F_(2, 16)_ = 0.4698 and p = 0.6335 for treatment effects), SOX2/p75-2-treated mice showed a significantly greater number of voiding contractions (Fig. 3F; F_(2, 16)_ = 11.02 and ***p = 0.0010 for treatment effects; **p = 0.0016 for SOX2/p75-2 vs. GFP, **p = 0.0037 for SOX2/p75-2 vs. p75-2, and p = 0.8291 for p75-2 vs. GFP) and enhanced bladder voiding efficiency (Fig. 3G; F_(2, 16)_ = 18.60 and ****p < 0.0001 for treatment effects; ***p = 0.0001 for SOX2/p75-2 vs. GFP, ***p = 0.0003 for SOX2/p75-2 vs. p75-2, and p = 0.7412 for p75-2 vs. GFP). Together, these results suggest that SOX2-mediated reprogramming alleviates SCI-induced urinary retention in contused mice.

**Figure 2.**
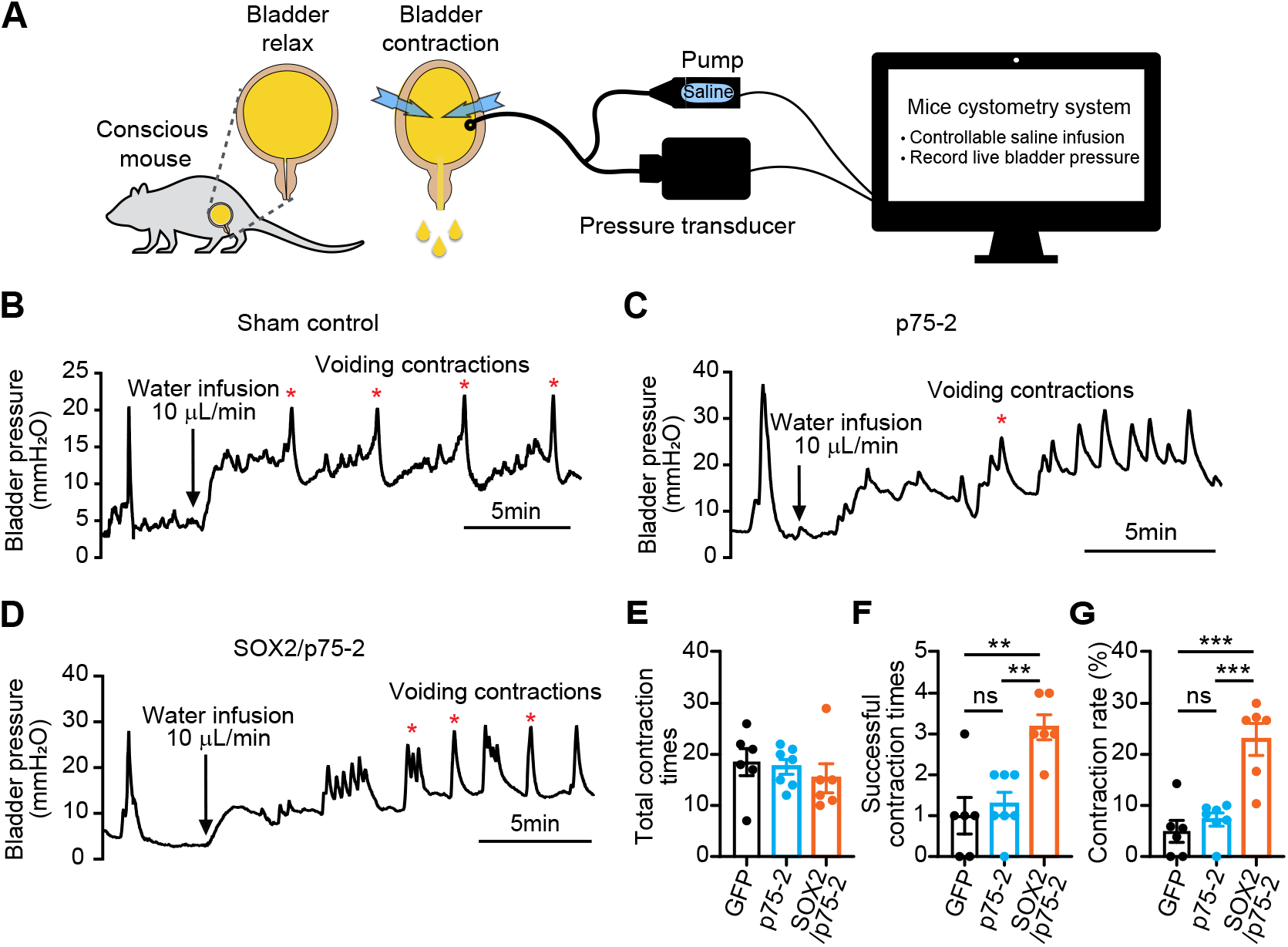
NG2 glia reprogramming reduces SCI-induced neurogenic bladder retention. (A) A schematic drawing of the procedure for cytometric evaluation of bladder function in the final week post SCI. (B-D) Histograms of bladder voiding contractions for the indicated experimental groups. (E) Quantification of contraction frequency (mean ± SEM, n = 6-7 mice per group; Ordinary one-way ANOVA: F_(2, 16)_ = 0.4698 and p = 0.6335 for treatment effects). (F) Quantification of successful contraction times (mean ± SEM, n = 6-7 mice per group; Ordinary one-way ANOVA: F_(2, 16)_ = 11.02 and ***p = 0.0010 for treatment effects; Tukey’s multiple comparisons test: **p = 0.0016 for SOX2/p75-2 vs. GFP, **p = 0.0037 for SOX2/p75-2 vs. p75-2; ns, not significant). (G) Quantification of contraction rate (mean ± SEM, n = 6-7 mice per group; Ordinary one-way ANOVA: F_(2, 16)_ = 18.60 and ****p < 0.0001 for treatment effects; Tukey’s multiple comparisons test: ***p = 0.0001 for SOX2/p75-2 vs. GFP, ***p = 0.0003 for SOX2/p75-2 vs. p75-2; ns, not significant).

**Figure 3.**
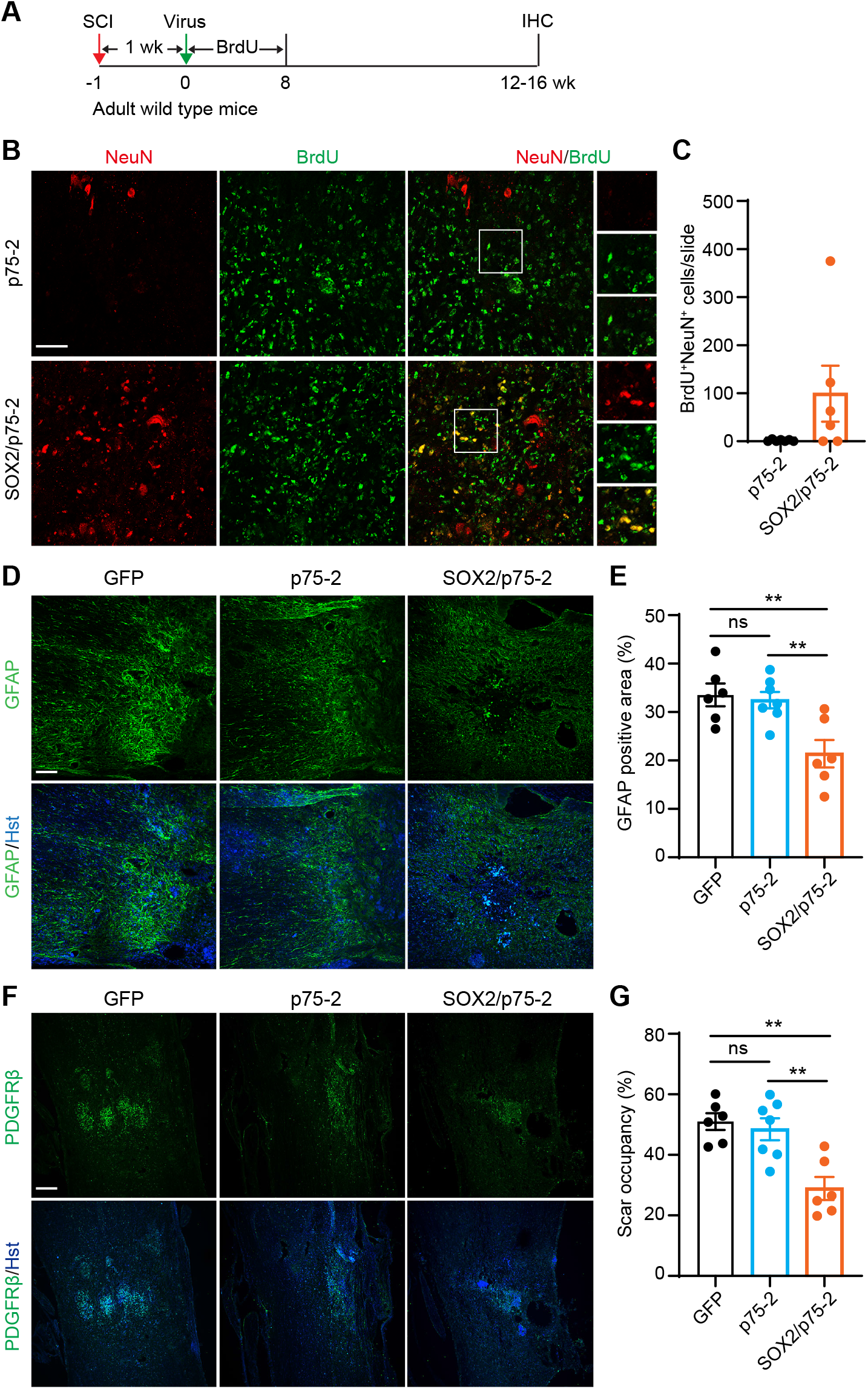
NG2 glia reprogramming induces adult neurogenesis and suppresses scarring. (A) Experimental scheme. Immunohistochemistry (IHC) was performed following completion of behavioral analyses. (B) Confocal images of BrdU-labeled neurons. Scale bar, 50 μm. (C) Quantification of BrdU-labeled new neurons (mean ± SEM; n = 6–7 mice per group). (D) Confocal images of GFAP expression surrounding the scar area. Nuclei were counterstained with Hoechst 33342 (Hst). Scale bar, 50 μm. (E) Quantification of GFAP-positive area (mean ± SEM, n = 6-7 mice per group; Ordinary one-way ANOVA: F_(2, 16)_ = 8.364 and **p = 0.0033 for treatment effects; Tukey’s multiple comparisons test: **p = 0.0057 for SOX2/p75-2 vs. GFP, **p = 0.0086 for SOX2/p75-2 vs. p75-2; ns, not significant). (F) Confocal images of PDGFRβ expression surrounding the scar area. Scale bar, 50 μm. (G) Quantification of scar occupancy (mean ± SEM, n = 6-7 mice per group, Ordinary one-way ANOVA: F_(2, 16)_ = 11.74and ***p = 0.0007 for treatment effects; Tukey’s multiple comparisons test: **p = 0.0013 for SOX2/p75-2 vs. GFP, **p = 0.0026 for SOX2/p75-2 vs. p75-2; ns, not significant).

### NG2 glia reprogramming induces adult neurogenesis and suppresses glial scarring

Following bladder function assessment, spinal cords were harvested for immunohistochemical analysis (Fig. 3A). Newly generated neurons were identified as BrdU-labeled NeuN^+^ cells. Although the number of new neurons varied among individual mice, substantial neuronal labeling was observed exclusively in the SOX2/p75-2-treated group, with no detection in mice receiving p75-2 alone (Fig. 3B, C).

Glial scar boundaries were assessed using GFAP immunostaining, which revealed a significant reduction in GFAP^+^ area in the SOX2/p75-2 group compared to either GFP or p75-2 controls (Fig. 3D, E; F_(2, 16)_ = 8.364 and **p = 0.0033 for treatment effects; **p = 0.0057 for SOX2/p75-2 vs. GFP, **p = 0.0086 for SOX2/p75-2 vs. p75-2, p = 0.9388 for p75-2 vs. GFP). Scar occupancy was further evaluated by PDGFRβ staining, showing a marked decrease in the SOX2/p75-2 group when compared to the controls (Fig. 3F, G; F_(2, 16)_ = 11.74 and ***p = 0.0007 for treatment effects; **p = 0.0013 for SOX2/p75-2 vs. GFP, **p = 0.0026 for SOX2/p75-2 vs. p75-2, p = 0.8604 for p75-2 vs. GFP). Together, these findings demonstrate that SOX2-mediated reprogramming of NG2 glia induces adult neurogenesis and attenuates glial scarring in the injured spinal cord.

### Significant correlation of neurogenesis and scar reduction to bladder function recovery

To assess the relationship between NG2 glial reprogramming and bladder function recovery, Spearman correlation analyses were conducted (Fig. 4A). The results revealed a significant positive correlation between improved urinary function and adult neurogenesis, as indicated by the number of BrdU^+^NeuN^+^ cells (Fig. 4A, B; R = 0.67, *p = 0.0167). Neurogenesis was also strongly negatively correlated with glial scarring, as measured by GFAP intensity (R = –0.81, **p = 0.0015) and glial scar occupancy (R = –0.76, **p = 0.0042), suggesting that reprogramming significantly reduces scarring. Conversely, increased glial scarring was associated with impaired urinary performance, with urine flow rate negatively correlating with both GFAP intensity (R = –0.59, *p = 0.0415) and glial scar occupancy (R = –0.59, *p = 0.0446). A strong positive correlation between GFAP intensity and glial scar area (R = 0.97, ***p = 1.3 × 10^−7^) further validated the consistency between these two scarring metrics. Together, these findings support a conclusion that SOX2-mediated reprogramming of NG2 glia induces neurogenesis, reduces glial scarring, and facilitates functional recovery of bladder control following spinal cord injury.

**Figure 4.**
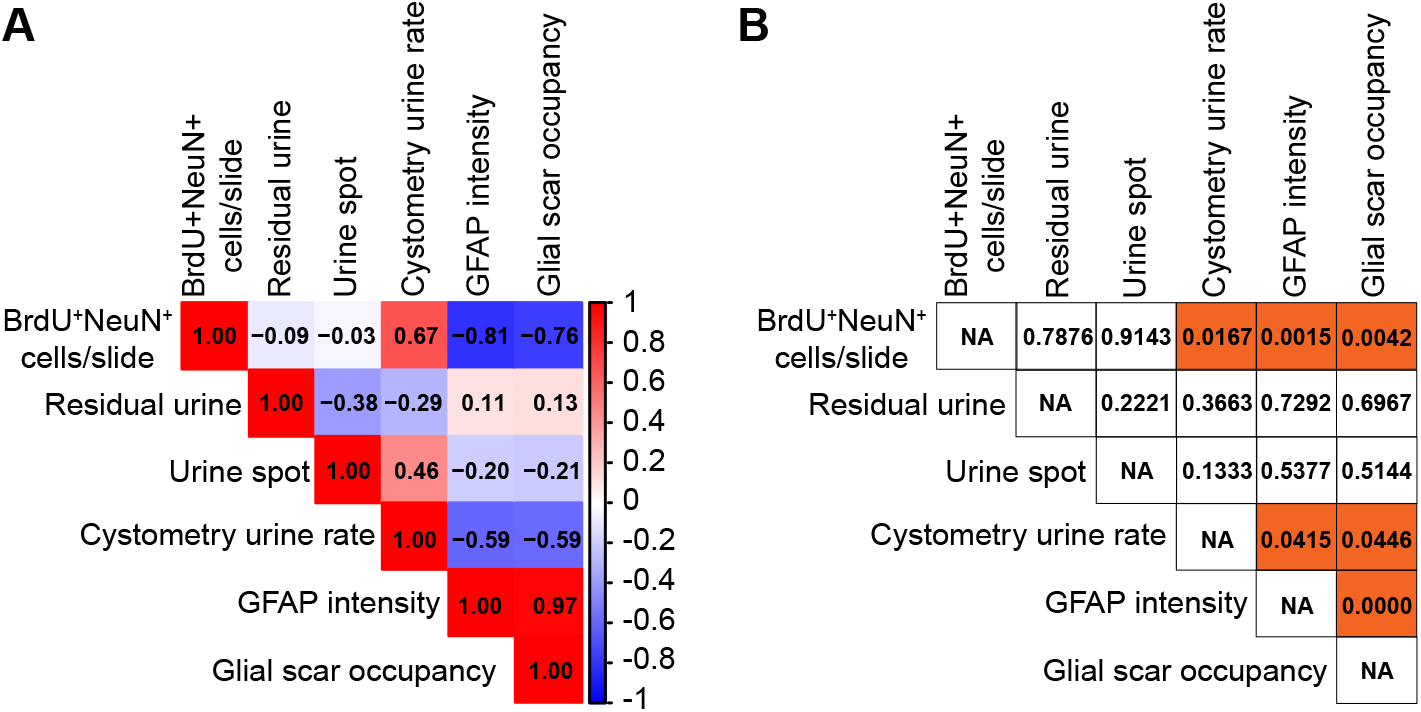
Spearman correlation of neurogenesis and scar reduction to bladder function recovery. (A) Spearman correlation matrix displaying the relationships between the indicated parameters. Corresponding R values are shown for each pairwise correlation. (B) P-value matrix for Spearman correlations showing the statistical values of each pairwise correlation. Corresponding p-values are indicated for each comparison.

## DISCUSSION

Neurogenic bladder remains one of the most challenging complications of SCI. Our findings demonstrate that SOX2-mediated in vivo reprogramming of NG2 glia significantly improves bladder control following contusive SCI. During a 2-hour observation period, SOX2-treated mice produced markedly larger urine spots compared to controls, indicating enhanced spontaneous voiding. To confirm that this reflected improved bladder function, we performed conscious cystometry, which revealed more efficient, high-amplitude voiding contractions in SOX2-treated mice relative to control groups. These results provide the first evidence that in vivo glial reprogramming may hold therapeutic potential for neurogenic bladder after SCI.

While recent advances in SCI research have led to improvements in locomotor and hand function ^9,18-20^, sensory and autonomic dysfunction--particularly neurogenic bladder--remains inadequately addressed. Notably, restoration of bladder function is consistently ranked as a top priority by individuals living with SCI ^21^. Although epidural stimulation can transiently enhance voiding ^22,23^, its benefits are lost once stimulation stops, and the underlying mechanisms remain poorly understood. In line with findings from cell transplantation studies ^24^, our results show that in vivo glial reprogramming can restore urinary function following severe contusive SCI. Unlike stem cell grafts, the reprogramming strategy utilizes the patient’s own cells, thereby avoiding the risks of immune rejection and tumorigenesis.

SOX2-mediated reprogramming of NG2 glia not only promoted the generation of new neurons within the injured T13 spinal cord but also significantly reduced glial scarring. These findings are consistent with our previous observations in the cervical hemisection model of SCI ^13^. Spearman correlation analysis revealed that bladder function recovery was positively associated with the number of newly induced neurons and negatively associated with the extent of glial scarring. These correlations raise important mechanistic questions regarding which aspect of reprogramming--neuronal replenishment, scar reduction, or both--primarily drives functional recovery. Moreover, it remains unclear whether the newly generated neurons contribute to bladder control by directly integrating into the autonomic circuitry or by indirectly modulating local spinal networks. Given prior evidence that reprogrammed neurons can form synaptic connections with both long-range and local circuits ^13^, both possibilities are plausible and warrant further investigation using synaptic tracing, chemogenetics, and electrophysiological approaches.

In summary, our findings demonstrate that SOX2-mediated in vivo reprogramming of NG2 glia improves bladder function after SCI by inducing neurogenesis and attenuating glial scarring. This study advances the concept of glial reprogramming as a regenerative strategy for neural repair and highlights its therapeutic potential for treating neurogenic bladder.

## MATERIALS AND METHODS

### Animals

The following mice were purchased from the Jackson Laboratory and maintained at local animal facility: wild-type C57BL/6J (stock #000664), R26R-tdT (Ai14, stock #007914) ^25^, Ascl1-CreER^T2^ (stock #012882) ^26^, Aldh1l1-CreER^T2^ (stock #031008) ^26^, and Pdgfra-CreER^T2^ (stock #032770) ^27^. Both adult male and female mice at 2 months of age and older were used for all experiments unless otherwise stated. All mice were housed under a controlled temperature and a 12-h light/dark cycle with free access to water and food in the animal facility. Sample sizes were empirically determined. Animal procedures and protocols were approved by the Institutional Animal Care and Use Committee at UT Southwestern or University of Texas Medical Branch.

### Spinal cord injuries and lentivirus injection

Adult mice were anesthetized with a cocktail of ketamine and xylazine. A laminectomy was performed at the indicated spinal segments. A contusion injury was produced by using the IH impactor (Precision Systems and Instrumentation, Lexington, KY) with a 1-mm tip and a force of 70 kdynes. After surgery, animals returned to their home cages and received manual bladder expressions twice daily. Lentivirus generation and titer determination were prepared as previously described^28,29^. For cellular and histological experiments, 1.5 µL of lentivirus (0.5-2 × 10^9^ pfu/mL) was manually injected into the spinal parenchyma using a Hamilton syringe fitted with a 34-gauge, 18-degree-beveled needle. For bladder function analysis in the T13 contusion injury model, viruses were delivered stereotaxically using a glass micropipette connected to a Nanoject III (Drummond) injector. Micropipettes were pulled to a tip diameter of ∼20-30 μm and mounted on the injector for precise delivery at 2 nL/min. A total of four stereotaxic injections were performed--two at 0.5 mm rostral and two at 0.5 mm caudal to the contusion center--to target glial cells at the lesion margins. At each site, bilateral injections (0.5 µL per site) were made at mediolateral (ML) ±0.5 mm from the midline and dorsoventral (DV) depths of 0.4 mm and 0.8 mm from the dorsal surface of the spinal cord.

### Tamoxifen and BrdU administration

Tamoxifen (Cayman chemical; Item No. 13258) was dissolved in a mixture of ethanol and sesame oil (1:9 by volume) at a concentration of 40 mg/mL and injected intraperitoneally at a daily dose of 1 mg/10g body weight for 5-7 days. Proliferating cells were labeled in vivo through administration with BrdU (Thermo Scientific Chemicals; Cat# H27260.06; 0.5g/L) in drinking water for the indicated durations.

### Immunohistochemistry

Mice were sacrificed with CO_2_ overdose and sequentially perfused with ice-cold phosphate-buffered saline (PBS) and 4% (w/v) paraformaldehyde (PFA) in PBS. Spinal cords were carefully dissected out, post-fixed overnight with 4% PFA at 4°C, and dehydrated in 30% sucrose for 2 days. Serial horizontal cryostat-sections (thickness: 17 µm) segment of the spinal cord spanning the injection/injury sites were collected on a cryostat (Leica). The immunostaining procedure was conducted as previously described ^29^. Spinal cord sections were stained with primary antibodies in blocking buffer (3% goat serum or 5% donkey serum in 0.3% PBST) for at least 18h at 4°C. The primary antibodies were rabbit anti-NeuN (1:1000, Abcam, Cat# ab177487), rat anti-BrdU (1:300, Bio-Rad, Cat# OBT0030), rabbit anti-DsRed (1:1000, Takara Bio, Cat# 632496), Goat anti-DCX (1:500, Santa Cruz Biotechnology, Cat# sc-8066), Guinea Pig anti-ASCL1 (1:100, Jane E. Johnson lab, UT Southwestern Medical Center), rabbit anti-PDGFRβ (1:1000, abcam, Cat# ab32570) and mouse anti-GFAP (1:1000, sigma, G3893). Detection was accomplished by incubation with Alexa Fluor- or Cy3-conjugated secondary antibodies (Life Technologies/Abcam) diluted in blocking buffer for 2 h at room temperature. Nuclei were counterstained with Hoechst 33342 (Hst; ThermoFisher, H1399). Images were captured by using the Zeiss LSM700, NIKON A1R or NIKON AXR confocal microscope. Confocal images were taken with a 20x objective, and the ImageJ program was used for cell counts.

### Scar occupancy

Scar core occupancy was quantified based on PDGFRβ expression, as previously described ^30-32^. Using Fiji (ImageJ), the total lesion area (Area_total) was defined by manually outlining the lesion boundary based on nucleus Hst counterstaining and morphological landmarks. PDGFRβ-positive regions were identified by manually thresholding the fluorescence channel and converting the image to binary format. To reduce background noise, objects smaller than 50 μm^2^ were excluded. The total PDGFRβ-positive area within the lesion (Area_PDGFRβ) was then calculated using the “Analyze Particles” function. Scar occupancy was expressed as the percentage of the lesion area occupied by PDGFRβ-positive staining using the following formula: Scar occupancy (%) = (Area_PDGFRβ / Area_total) × 100%.

### GFAP-positive area

This was quantified as previously described with modifications ^32-35^. GFAP-stained images were converted to 8-bit grayscale using ImageJ. For each section, a threshold was determined based on GFAP staining intensity in a region remote from the lesion core. GFAP-positive signal, representing astrogliosis, was then thresholded within a 500 μm region both rostral and caudal to the lesion center. The GFAP-positive area was expressed as a percentage of the total area analyzed.

### Voided stain on paper (VSOP) analysis

This procedure was reported previously ^24^ with modifications. Mice were individually placed in 30 × 15 cm cages on sheets of filter paper (Whatman) covering the whole cage floor area to measure the voiding pattern. Prior to VSOP tests, all mice were habituated for 10 minutes followed by bladders being manually expressed (to empty the bladder), and 50 μL/g body weight of saline was injected subcutaneously into all tested mice. All tested mice were placed in the cage for 120 minutes, and filter paper was collected at the end of experiments. Collected filter papers were stained with Ninhydrin (Sigma) and digitally scanned. The color images were analyzed by Image J (NIH). The diameter of each urine spot was measured (overlapping urine spots were counted as one). Only the urine spots with a diameter bigger than 3.5 cm were counted, which means the urine volume is more than 100 µL. The number and diameter of urine spots were quantified.

### Catheter implantation in the bladder

All procedures used have been detailed previously ^36,37^. All tested mice were first habituated in a Small Animal Cystometry Lab station (Catamount Research and Development) in a recording cage for 30-40 mins on two consecutive days before catheter implantation. On the surgery day, mice were placed on a heated pad and anesthetized with ketamine/xylazine (100/10 mg/kg). Each bladder dome was exposed and exteriorized through a middle abdominal incision. A 7-cm PE10 tube was used as the catheter and inserted into the bladder. A loose tie of monofilament suture (6-0) was used to secure the catheter in the bladder. Then the end of the catheter was pulled out through an incision at the back of the mouse’s neck. The abdominal wound and neck incision were closed, and each mouse was allowed to recover for at least 48 hours prior to cystometry recordings. After each surgical procedure, 1 mL of saline was injected subcutaneously. Postoperative care was comprised of manual bladder compression and animal activity was monitored twice daily.

### Conscious cystometry

This procedure was reported previously ^24^ with modification. At 2 days post-catheter implantation, conscious csytometry was performed. The tubing was connected to one port of a pressure transducer; the other port of the pressure transducer was connected to a syringe pump, which allowed room-temperature saline to be continuously infused at a rate of 10 µL/min. In intact mice, we observed 4-5 reproducible micturition cycles that were recorded after an initial stabilization period of 30 mins. Therefore, all the animals were tested in 30 mins. To confirm the bladder contraction can be recorded in the conscious mice, the bladder was manually expressed in all tested mice prior to saline infusion and the bladder pressure changes were observed in all tested mice. Assessment of bladder function was unbiased since the cystometric parameters were measured using an automated cystometry analysis software SOF-552 (Catamount Research and Development). These included maximal voiding pressure (cm H_2_O), the number of voiding and non-voiding urinary bladder contractions per 30 min. Non-voiding contractions were defined as rhythmic intravesical pressure rise (>5 cm H_2_O from baseline pressure) without a release of fluid from the urethra. At the completion of conscious cystometry, the mice were humanely euthanized, spinal cord and bladder tissues were collected for histology.

### Statistical analysis

Data are presented as mean ± SEM. Ordinary one-way ANOVA and Tukey’s post hoc multiple comparisons were used for both histological and bladder function analysis. Spearman correlation analysis was performed using R software. All analyses were conducted blind to experimental conditions. A p value < 0.05 was considered significant. Significant differences are indicated by *p < 0.05, **p < 0.01, ***p < 0.001, and ****p < 0.0001.

## AUTHOR CONTRIBUTIONS

W.T., J.S., P.Z., W. L., B.C., C.L.Z. conceived and designed the experiments. W.T., J.S., P.Z., W. L., performed the experiments. T.S., X.Z. provided critical reagents and scientific inputs. Y.Z. maintained mouse colonies. W.T., J.S., P.Z., W. L., B.C., C.L.Z. analyzed data. W.T., B.C., C.L.Z. wrote the manuscript. All authors reviewed and approved the manuscript.

## ACKNOWLEDGEMENTS

We thank members of the Zhang laboratory for discussions and reagents and J. E. Johnson (UT Southwestern, USA) for ASCL1 antibody. C.L.Z. is a W. W. Caruth, Jr. Scholar in Biomedical Research. The work in Zhang laboratory was supported by NIH (NS127375, NS117065, NS131489, NS111776), and the work in Chen laboratory was supported by the Mission Connect, TIRR Foundation, Criag Neilson Foundation, and UTMB Claire E. Hulsebosch Chair in Neurological Recovery endowment.

