## Supplemental Figures for "in vivo Reprogramming of NG2 Glia Improves Bladder Function After Spinal Cord Injury"

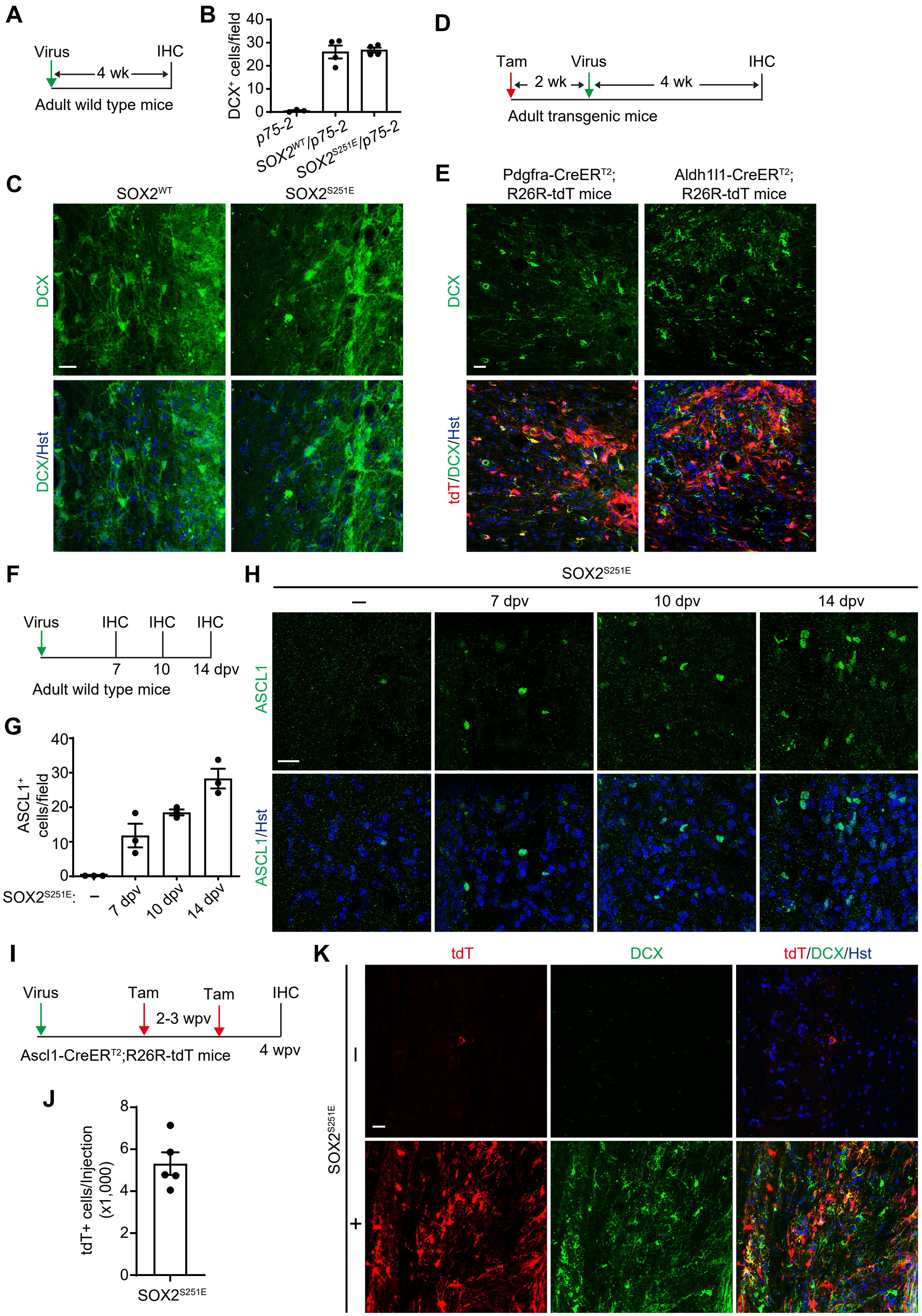
A
Virus
IHC
4 wk
Adult wild type mice
B
DCX+ cells/field
p75-2
SOX2WT/p75-2
SOX2S251E/p75-2
C
SOX2WT
SOX2S251E
DCX
DCX/Hst
D
Tam
Virus
IHC
2 wk
4 wk
Adult transgenic mice
E
Pdgfra-CreERT2; R26R-tdT mice
Aldh1l1-CreERT2; R26R-tdT mice
DCX
tdT/DCX/Hst
F
Virus
IHC
IHC
IHC
7
10
14 dpv
Adult wild type mice
G
ASCL1+ cells/field
SOX2S251E:
–
7 dpv
10 dpv
14 dpv
H
SOX2S251E
–
7 dpv
10 dpv
14 dpv
ASCL1
ASCL1/Hst
I
Virus
Tam
Tam
IHC
2-3 wpv
4 wpv
Ascl1-CreERT2;R26R-tdT mice
J
tdT+ cells/Injection (x1,000)
SOX2S251E
K
tdT
DCX
tdT/DCX/Hst
SOX2S251E
–
+

**Supplementary Figure S1. SOX2-mediated glial reprogramming in the adult mouse spinal cord.**

(A) Experimental scheme. SOX2-expressing lentiviruses were injected into the adult spinal cord, and tissues were analyzed 4 weeks (wk) later using immunohistochemistry (IHC).

(B) Quantification of SOX2-induced $DCX^+$ immature neurons (mean ± SEM, n = 3-4 mice per group). Notably, $DCX^+$ cells were absent in the p75-2–only condition.

(C) Confocal images of $DCX^+$ cells induced by either wild-type SOX2 ($SOX2^{WT}$) or the phospho-mimetic mutant SOX2 ($SOX2^{S251E}$). Scale bars, 20 μm.

(D) Experimental scheme for tracing the cellular origin of mutant SOX2-reprogrammed cells. Adult transgenic mice were treated with tamoxifen (Tam), followed by intraspinal viral injections, and analyzed 4 weeks later.

(E) Confocal images showing that NG2 glia are the cellular origin of mutant SOX2-reprogrammed cells in the adult spinal cord. NG2 glia and astrocytes were lineage-traced using *Pdgfra-CreER$^{T2}$;R26R-tdT* and *Aldh1l1-CreER$^{T2}$;R26R-tdT* mice, respectively. Scale bar, 20 μm.

(F) Experimental scheme for a time-course analysis of mutant SOX2-mediated reprogramming. dpv, days post virus-injection.

(G) Quantification of mutant SOX2-induced $ASCL1^+$ neural progenitors.

(H) Confocal images showing induction of $ASCL1^+$ neural progenitors by mutant SOX2. Scale bar, 20 μm.

(I) Experimental scheme for tracing the progeny of mutant SOX2-induced $ASCL1^+$ progenitors in the adult spinal cord. wpv, weeks post virus-injection.

(J) Quantification of mutant SOX2-induced $tdT^+$ cells at 4 wpv (mean ± SEM, n = 5 mice).

(K) Confocal images showing robust detection of $DCX^+$tdT+ cells in mutant SOX2-expressing spinal cord. Scale bar, 20 μm.
